# Rapid De Novo Antibody Design with GeoFlow-V3

**DOI:** 10.1101/2025.10.20.682964

**Authors:** BioGeometry Team, Jian Tang

## Abstract

Recent years have witnessed striking advances in miniprotein design, yet *de novo* antibody discovery remains challenging, marked by low binding rates and the need for extensive, labor-intensive experimental screening of millions of candidates. This technical report introduces GeoFlow-V3, a unified atomic generative model for structure prediction and protein design. GeoFlow-V3 delivers improved accuracy on antibody–antigen complex structure prediction relative to our previous version, and its performance is further enhanced when experimental constraints or prior knowledge are provided, enabling precise control over both folding and design. The model also demonstrates reliable ability to discriminate binders from non-binders based on its confidence scores. Leveraging this capability, we build a GeoFlow-V3 *in silico* pipeline to design no more than 50 nanobodies per therapeutically relevant target *de novo*, completing a single round of wet-lab characterization in under three weeks. GeoFlow-V3 identifies at least one binder for 8 tested epitopes and achieves an average hit rate of 15.5%, representing a two-orders-of-magnitude improvement over prior computational pipelines. These results position GeoFlow-V3 as an appealing platform for rapid, AI-driven therapeutic antibody discovery, significantly reducing experimental screening demands and offering a powerful avenue to tackle previously undruggable targets. A demo of GeoFlow-V3 can be accessed via <u>prot.design</u> for non-commercial use.

## 1. Introduction

Proteins are the molecular workhorses of life, executing nearly every cellular function, from catalyzing metabolic reactions to transmitting signals and maintaining structural integrity. Among these, antibodies play a central role in immune defense, recognizing and neutralizing a vast array of pathogens with exquisite specificity. Their unique ability to bind diverse antigens has made antibodies indispensable not only to human physiology but also to modern medicine (Buss et al., 2012; Lu et al., 2020). Over the past three decades, monoclonal antibodies and their derivatives have transformed the therapeutic landscape, enabling targeted treatments for cancer, autoimmune disorders, infectious diseases, and more (Zahavi and Weiner, 2020).

While machine learning has delivered remarkable advances in biomolecular structure prediction (Abramson et al., 2024; Baek et al., 2023; Corley et al., 2025; Jumper et al., 2021) and in the *de novo* design of compact miniprotein binders (Cao et al., 2022; Pacesa et al., 2025; Ren et al., 2025; Watson et al., 2023; Zambaldi et al., 2024), *de novo* antibody discovery remains a fundamentally harder problem. Antibody–antigen interfaces are dominated by conformationally flexible complementarity-determining region (CDR) loops, which are far more difficult to model than the relatively rigid *α*-helical scaffolds typical of miniproteins. Current structure-prediction models struggle to capture antibody–antigen complexes with sufficient accuracy, and robust virtual screening or affinity ranking among candidate binders remains elusive. Although several computational pipelines for *de novo* antibody generation have been proposed recently (Bang et al., 2025; Bennett et al., 2024; Bio and Biswas, 2025), they still depend on labor-intensive high-throughput screening (e.g., phage or yeast display) that can take months. Moreover, oligonucleotide-pool–based assays limit the length of designable regions, making fully *de novo* antibody generation costly and time-consuming.

An ideal computational framework would be one that couples atomic-level structure prediction with generative design, discriminates true binders from non-binders accurately, and requires only minimal experimental validation. Contemporary efforts point in this direction, including Chai-2 (Chai-Discovery et al., 2025), a diffusion-based approach achieving double-digit hit rates with limited wet-lab testing, and Germinal (Mille-Fragoso et al., 2025), which integrates AlphaFold Multimer (Evans et al., 2021) gradients with an antibody language model (Shuai et al., 2023) to hallucinate target-binding antibodies.

In this technical report, we introduce GeoFlow-V3, a unified atomic diffusion model that not only integrates high-fidelity complex modeling with generative design, but also incorporates an *in silico* evolution procedure, enabling rapid, low-*N* antibody discovery. Our main contributions are:

- GeoFlow-V3 achieves high-accuracy docking and reliable binder discrimination. It improves DockQ scores for antibody–antigen complexes and, when augmented with experimental constraints or prior knowledge, allows precise control over folding and design. It robustly distinguishes correct complexes from decoys and identifies binders from non-binders based on confidence scores.
- GeoFlow-V3 implements an *in silico* evolution strategy that enhances candidate quality. This strategy increases the number of designs passing structural filters and raises maximal achievable confidence scores.
- GeoFlow-V3 enables rapid, low-*N* antibody discovery. We design and experimentally test up to 50 nanobodies per campaign for 8 campaigns spanning 5 therapeutically relevant targets (including 3 multi-epitope targets). GeoFlow-V3 identifies at least one confirmed binder per campaign, achieves a double-digit average hit rate—over two orders of magnitude higher than prior computational pipelines, and generates novel nanobodies with nanomolar binding affinity.

## 2. Methods

### 2.1. A unified atomic diffusion model for structure prediction and *de novo* design

GeoFlow-V3 is a unified atomic diffusion model that integrates protein structure prediction and *de novo* design across diverse biological modalities. GeoFlow-V3 is trained on high-quality predicted structures and the PDB (Berman et al., 2000), with a training data cutoff of 30 June 2024. Building on the *pseudo protein sequence* mechanism introduced in GeoFlow-V2 (BioGeometry, 2025), the model accepts full sequences for structure prediction and partially or fully masked sequences for *de novo* design.

Regarding antibody–antigen complexes, GeoFlow-V3 increases the fraction of predictions achieving high DockQ accuracy—a stringent but critical metric for *de novo* antibody design—by 45% compared to the previous generation, GeoFlow-V2. The model’s confidence scores effectively discriminate binding potential, with high-confidence predictions exhibiting over 80% precision in distinguishing correct from incorrect structures. In addition, GeoFlow-V3 enables zero-shot, target structure-conditioned antibody design, generating binders toward user-specified binding sites. These sites can be defined either as precise hotspot residues or as a set of coarse-grained binding residues, allowing flexible design without requiring an initial binder for accurate epitope selection. For particularly challenging targets, GeoFlow-V3 leverage test-time scaling (Ma et al., 2025) to simulate the evolution of antibodies, iteratively refining candidate designs guided by confidence scores.

### 2.2. Simulating evolution of antibodies with GeoFlow-V3

Antibody diversity arises from V(D)J recombination, somatic hypermutation, and in vivo affinity maturation (Chi et al., 2020), collectively generating a broad repertoire capable of recognizing diverse antigens. In contrast, previous versions of GeoFlow adopt a one-round “generation–filtering” pipeline, thus often struggle to identify true binders within a limited number of assayed designs, particularly when starting from noisy epitopes or uncertain antigen structures.

Inspired by natural antibody evolution and recent advances in test-time scaling of diffusion models (Ma et al., 2025; Uehara et al., 2025), we hypothesize that GeoFlow-V3 can be steered toward multi-round *de novo* antibody design, by leveraging its constraint-conditioning capability (Section 3.1.3). Starting from initial promising candidates, we aim to iteratively refine both sequences and structures through a partial diffusion mechanism (Glögl et al., 2024), which initializes the denoising process with a partially noised input, thereby biasing the model toward desired interaction patterns.

For **structure redesign**, we apply the reverse noising process to the clean structure for 35%–45% of the diffusion timesteps before invoking the model to denoise it. For **sequence redesign**, we explore two complementary strategies: (1) **Full CDR redesign**, in which all CDR residues are redesigned jointly with the structure; and (2) **Partial CDR redesign**, in which we identify suboptimal residues based on a residue-level **CDR-ipAE** (Section 3.2.1), defined as the mean PAE of each CDR residue with respect to the target, and selectively redesign those exceeding a predefined threshold.

By iteratively refining promising candidates in this manner, guided by model-assigned confidence scores or external reward functions (Figure 2), GeoFlow-V3 effectively traverses the sequence–structure landscape in a process analogous to natural affinity maturation, ultimately yielding high-confidence binders suitable for downstream experimental validation (Section 3.3).

**Figure 1.**
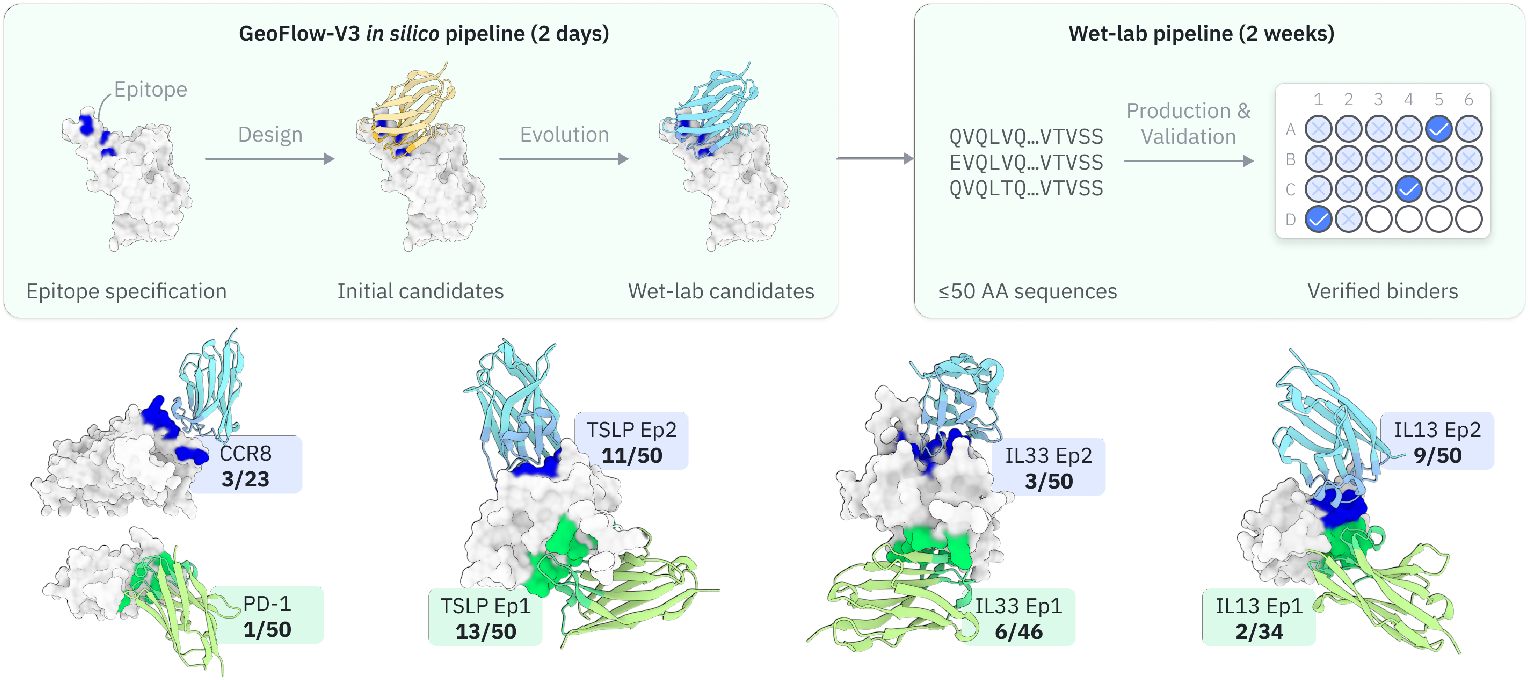
From epitope to verified binders within 3 weeks. **(Top)** The GeoFlow-V3 *de novo* antibody design pipeline. Given a specified epitope, GeoFlow-V3 first generates target-specific antibody binders, then perform *in silico* evolution to enhance their bind rate. The top-ranked (no more than 50) designs are then advanced to wet-lab validation, with single-concentration BLI for binder identification and multi-concentration BLI for binding affinity characterization. **(Bottom)** Structure of identified binders on 8 distinct, therapeutically-relevant epitopes. GeoFlow-V3 achieves an average hit rate of 15.5% across multiple targets. Target names, number of identified binders and number of tested binders are shown next to the structure.

**Figure 2.**
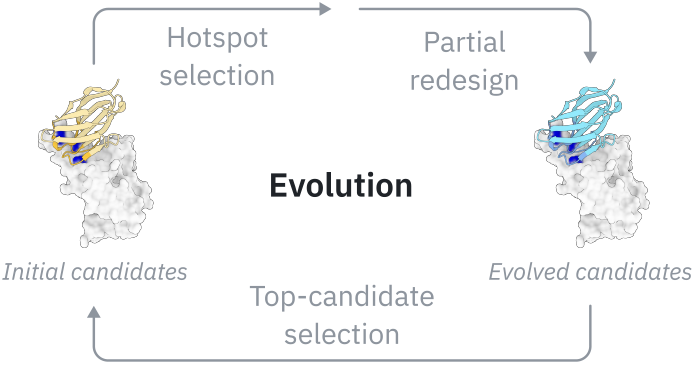
Illustration of the *in silico* antibody evolution process. Starting from initial candidates, GeoFlow-V3 partially redesigns both sequences and structures, progressively evolving them toward higher-confidence candidates.

### 2.3. Experimental validation of design capabilities on diverse targets

To evaluate the zero-shot antibody design capability of GeoFlow-V3, we experimentally validate its performance on *de novo* VHH design tasks. We select 5 therapeutically relevant antigens with known three-dimensional structures, most of which lack reported antigen:VHH structure in the Protein Data Bank (PDB). For each design campaign, we identify up to 10 structurally meaningful epitope residues from ligand–receptor complex structures (Table S1). At each forward pass, GeoFlow-V3 randomly subsamples these residues to create input epitope constraints. To probe epitope-specific design, we define two distinct epitope sets for 3 of the 5 targets, resulting in 8 independent VHH design campaigns. No known VHH binder—if any exists—is used at any stage of the design loop.

As GeoFlow-V3 demonstrates improved high-fidelity antibody–antigen structure prediction and reliable binding discrimination, we hypothesize that the traditional time-consuming display-based screening (Boder et al., 2012; Hoogenboom et al., 1998) could be omitted. Instead, we opt for **a single round of wet-lab characterization via direct gene synthesis**. For each VHH campaign, we select up to 50 generated designs for experimental validation. Binding strength is assessed by bio-layer interferometry (BLI). Following Chai-2’s practice (Chai-Discovery et al., 2025), positive binders are defined as designs exhibiting a binding-positive curve signature with a signal both greater than 0.1 nm above background and at least 300% of the background response. We provide further details for these settings in the following sections.

## 3. *In silico* Experiments

We perform *in silico* benchmarking of GeoFlow-V3 across three evaluation settings: (1) antibody–antigen structure prediction on a curated low-homology dataset; (2) binder discrimination, assessing the ability to distinguish true binders from non-binders and to correlate model confidence scores with experimental binding affinities; and (3) case studies demonstrating the effectiveness of *in silico* antibody evolution.

### 3.1. Antibody-antigen structure prediction

#### 3.1.1. Setup

Our evaluation protocol largely follows that of GeoFlow-V2 (BioGeometry, 2025) with updated baselines and expanded metrics. We assess GeoFlow-V3 on a carefully curated low-homology test set, following AlphaFold 3’s interface-validation guidelines (Supplementary Section 5.8). To build the benchmark, we collect all PDB antibody–antigen complexes released between 30 June 2024 and 30 January 2025, crop antibodies to their Fv regions, and retain only complexes containing 256-1024 residues to maintain computational efficiency. This yields a final test set of 104 high-quality antibody–antigen complexes.

For comparison, we include recent state-of-the-art AlphaFold 3 reproduction methods: **Chai-1** (Chai-Discovery et al., 2024), **Protenix-v0.6.0** (Protenix et al., 2025), and **Boltz-2** (Wohlwend et al., 2024). We also benchmark against **AlphaFold Multimer V2.3** (Evans et al., 2021) and our previous **GeoFlow-V2** (BioGeometry, 2025). Diffusion-based approaches, including GeoFlow-V3, generate 50 candidate structures per target (10 random seeds × 5 diffusion samples). For AlphaFold Multimer V2.3, we ensemble five model checkpoints across ten random seeds with dropout enabled at inference. All predictions are scored with the DockQ metric (Basu and Wallner, 2016) under two settings: Top-1, the single highest-ranked prediction, and Top-10, the best among the ten highest-ranked predictions. We report success rates for three quality tiers—Acceptable, Medium, and High—defined by DockQ thresholds of 0.23, 0.49, and 0.80, respectively.

#### 3.1.2. GeoFlow-V3 achieves state-of-the-art performance on antibody-antigen docking

Figure 3A summarizes docking accuracy across all evaluated methods. Main findings are as follows.

**Figure 3.**
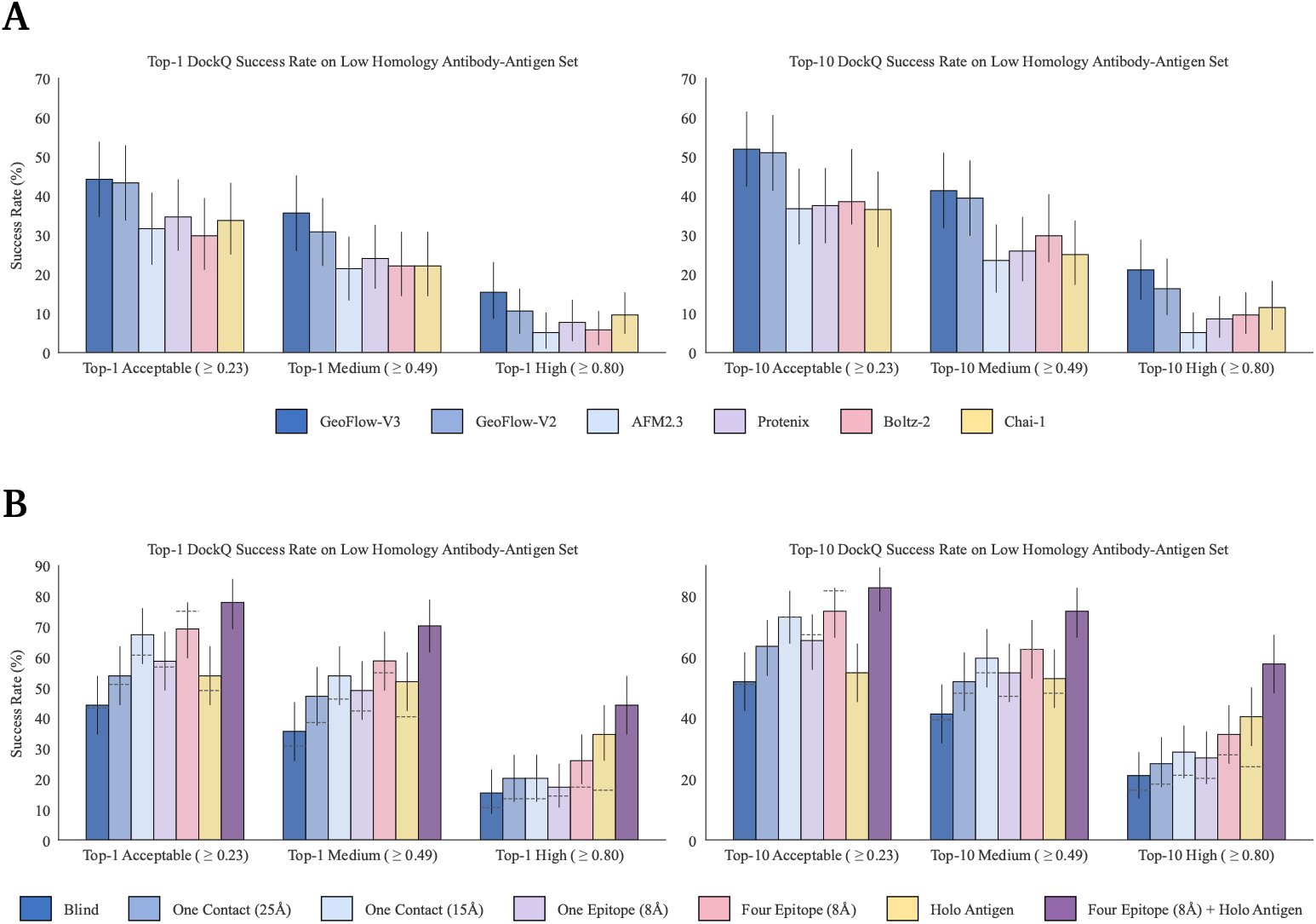
Antibody–antigen structure-prediction performance across evaluated models. Bar heights represent mean success rates over all test cases, and error bars denote 95% confidence intervals estimated from 10,000 bootstrap resamples. **(A)** DockQ-based comparison of antibody-antigen struture predictions for each model under two evaluation settings: *Top-1*, the highest-ranked prediction, and *Top-10*, the best prediction among the ten top-ranked candidates. Success rates correspond to the percentage of antibody–antigen complexes achieving DockQ scores above 0.23, 0.49, and 0.80, which define the Acceptable, Medium, and High quality tiers, respectively. **(B)** Benchmark of GeoFlow-V3 on antibody-antigen struture prediction under different constraint features, using the same metrics as in (A). Dashed lines indicate the corresponding performance of GeoFlow-V2, where available. *Blind*: Vanilla docking with sequence-only inputs. *One Contact (15Å / 25Å)*: A single antibody–antigen residue pair within 15Å or 25Å C*α* distance, provided to the model as a contact constraint. *One Epitope (8Å) / Four Epitopes (8Å)*: One (or four) antigen residues with C*α* atoms located within 8Å of the antibody, supplied as epitope constraints. *Holo Antigen*: Noise-perturbed bound antigen structures provided as conditioning features. *Four Epitopes (8Å) + Holo Antigen*: Combined setting with both holo antigen conditioning and four epitope constraints.

- **Clear performance gains over GeoFlow-V2**. GeoFlow-V3 delivers measurable improvements in antibody–antigen structure prediction relative to GeoFlow-V2. Although the increase in *Acceptable*-quality predictions is modest, GeoFlow-V3 substantially raises the rate of *High*-quality complexes, reaching a 15.4% Top-1 success rate and 19.2% Top-10 success rate, representing 45% (Top-1) and 18% (Top-10) relative gains over GeoFlow-V2.
- **High-DockQ as a proxy for design capability**. High-DockQ success usually demands sub-angstrom interface RMSD accuracy. We hypothesize that the High-DockQ rate of a model serves as a strong indicator of its *de novo* antibody-design capability. This is also hinted in the Chai-2 report (Chai-Discovery et al., 2025), which highlighted improved High-DockQ rate compared to Chai-1 (34% vs 17%, on a different test set). We note that despite recent advances, there is still considerable room for improvement in this metric.
- **Comparison to other foundation models**. Chai-1, Protenix-v0.6.0, and Boltz-2 are strong baselines on our benchmark, each achieving competitive results. Manual inspection reveals that different models succeed on partially disjoint subsets of targets, suggesting potential benefits from ensembling multiple foundation models in real-world applications. Surprisingly, AlphaFold Multimer V2.3 can still match diffusion-based methods when extensive sampling strategies are applied (Wallner, 2023).

#### 3.1.3. Constraints guiding enhances docking performance

In many practical settings, additional experimental or computational cues can guide structure prediction and de novo design. For antibody–antigen complexes, for example, epitope hints from wet-lab assays (Haynes et al., 2021; Moreira et al., 2007) or prior structural data can help focus the search space. Likewise, in *de novo* protein design, explicit target-structure conditioning or user-defined epitope constraints are often essential for steering the model toward a desired binding interface.

GeoFlow-V3 supports a broad range of constraint features, including:

- **Epitope Constraint**: user-specified residues or residue sets that the predicted (or designed) binder must contact, optionally subsampled during generation.
- **Contact Constraint**: pairwise residue–residue contacts or distance restraints that restrict the generated complex geometry, optionally subsampled during generation.
- **Target-Structure Constraint**: noise-perturbed distance maps extracted from *target* structures. Unlike the template features in AlphaFold, GeoFlow-V3 accommodates both intra-chain and inter-chain conditioning.
- **Initial-Guess Constraint**: inspired by the AlphaFold 2 initial guess strategy (Bennett et al., 2023), GeoFlow-V3 can seed the diffusion process with a user-provided starting conformation, which is a critical capability that enables test-time scaling to simulate antibody evolution.

In the following analyses, we evaluate how different constraint features enhance GeoFlow-V3’s docking accuracy on the low-homology antibody–antigen benchmark, using the same metrics as in the unconstrained setting. We compare seven inference modes: (1) **blind docking** with sequence-only inputs; (2 & 3) a **single contact** constraint specifying one antibody–antigen residue pair within either 15Å or 25Å C*α* distance; (4) a **single epitope** constraint selecting one random antigen residue within 8Å of the antibody; (5) **four epitope** constraints with four such antigen residues; (6) a **holo-antigen structure**-conditioning constraint that provides a noise-perturbed target structure while still requiring full complex prediction from random initialization; and (7) a **combined setting** that applies both holo-antigen conditioning and four epitope constraints.

Figure 3B summarizes the results, and the main findings are discussed below.

- Consistent with our earlier findings (BioGeometry, 2025), structural constraints substantially improve docking performance over the unconstrained baseline, underscoring their value when prior experimental data or design hotspots are available. GeoFlow-V3 yields stronger results than GeoFlow-V2 across most settings, with the largest gains in medium- and high-accuracy predictions.
- Among individual constraint types, holo-antigen conditioning provides unique advantages for high-accuracy predictions (DockQ *>* 0.80). This is especially relevant for practical applications, such as antibody optimization or antibody design, where bound antigen structures might be available. Four-epitope constraints deliver the best overall performance in the acceptable and medium-accuracy ranges, confirming the model’s ability to follow specified binding hotspots, a capability that is critical for *de novo* antibody design.
- Combining holo-antigen conditioning with four epitope constraints yields the strongest results across all metrics, reaching a 44.2% Top-1 and 57.7% Top-10 high-DockQ success rate. Although such detailed information is available in limited settings, these findings highlight the particular value of combining structural and epitope cues in antibody design and optimization tasks.

#### 3.1.4. Docking confidence scores distinguish correct from incorrect structures

Antibody–antigen structure prediction remains challenging despite recent advances in diffusion-based generative models for protein structure prediction. In practice, a large number of candidate complexes are often generated to increase structural diversity, after which only the highest-ranked predictions are selected (Abramson et al., 2024). It is therefore essential to evaluate a model’s ability to distinguish correct from incorrect predictions and to assess the reliability of its own outputs. A well-calibrated confidence measure is likewise crucial for high-quality data distillation (Jumper et al., 2021).

To investigate this capability, we analyze the top-ranked predictions for all 104 antibody–antigen complexes in our curated low-homology test set and examine the relationship between structural quality, measured by DockQ scores, and GeoFlow-V3’s confidence scores. Treating the confidence score as a binary classifier with a threshold of 0.8, we define predictions with confidence score *>* 0.8 and DockQ score *>* 0.23 as true positives. As shown in Figure 4, across different constraint settings, low-confidence predictions are rarely correct, whereas high-confidence predictions achieve over 80% precision in identifying Acceptable structures. These results highlight the value of confidence estimation for ranking candidates and for reliable data augmentation.

**Figure 4.**
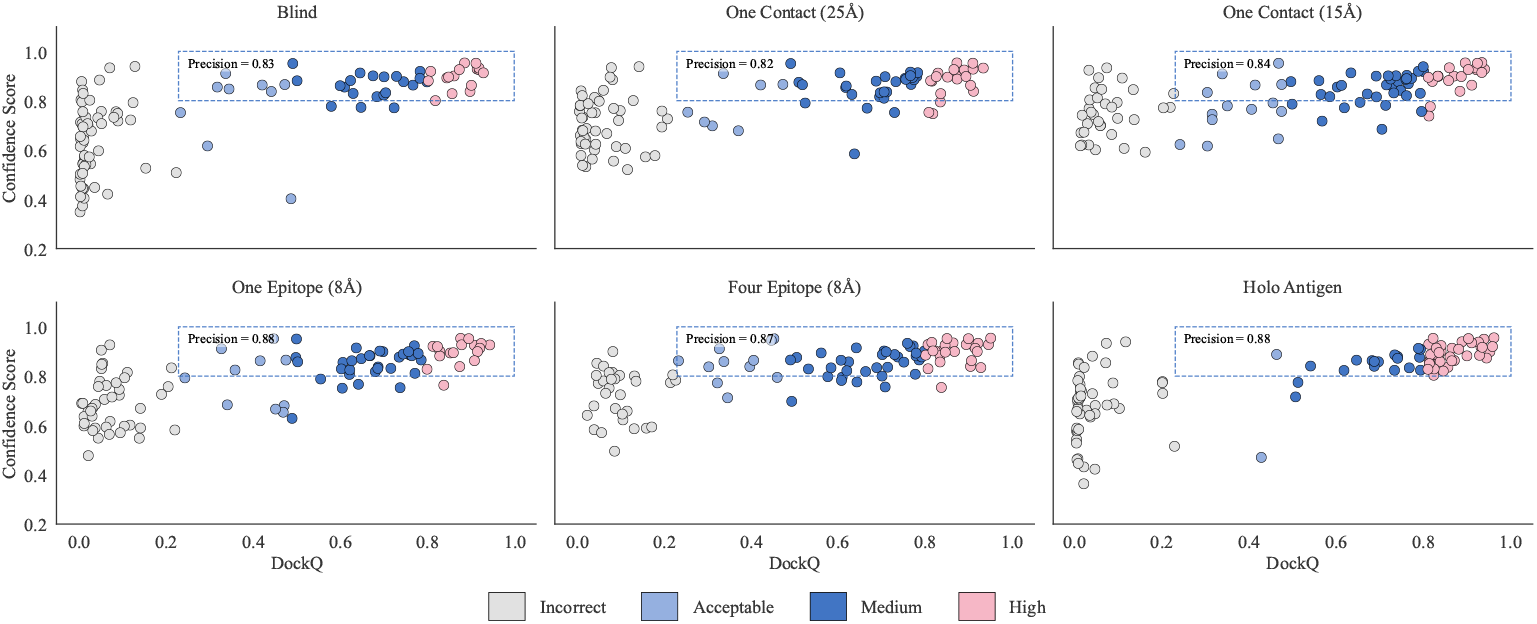
Visualization of DockQ scores versus GeoFlow-V3’s confidence scores for Top-1 predictions across 104 antibody–antigen complexes in the low-homology test set. The dashed box marks predictions with DockQ score *>* 0.23 and confidence score *>* 0.8, representing cases that meet both the Acceptable-quality threshold and the high-confidence criterion. The precision reported within this box corresponds to treating a confidence threshold of 0.8 as a binary classifier for correct predictions.

### 3.2. Binder identification performance

#### 3.2.1. Setup

While millions of candidate binders can be generated *in silico*, wet-lab validation remains a major bottleneck. Computational filtering and ranking are therefore essential for prioritizing the most promising designs before costly experimental testing. Here, we assess GeoFlow-V3’s ability to distinguish true binders from non-binders following the evaluation protocol of GeoFlow-V2, and we examine the correlation between model confidence scores and experimentally measured affinities using a subset of SKEMPI mutation data (Jankauskaitė et al., 2019).

We briefly summarize the ten datasets used for binder discrimination (full statistics in Table 1):

**Table 1.** Summary of binder/non-binder datasets used for discrimination evaluation.

|  | HER2<br>Trastuzumab | 5A12<br>VEGF | 5A12<br>ANG-2 | TSLP | FXI | IL36R | TNFRSF9 | C5 | ACVR2B | IL17A |
| --- | --- | --- | --- | --- | --- | --- | --- | --- | --- | --- |
| PDB ID | 1N8Z | 4ZFG | 4ZFF | 5J13 | 6HHC | 6U6U | 6A3W | 5I5K | 5NGV | 6PPG |
| Binders | 417 | 701 | 120 | 115 | 34 | 29 | 29 | 30 | 15 | 6 |
| Non-binders | 823 | 499 | 1,080 | 61 | 82 | 148 | 156 | 168 | 197 | 173 |
| Binder Rate (%) | 33.6 | 58.4 | 10.0 | 65.3 | 29.3 | 16.4 | 15.7 | 15.2 | 7.1 | 3.4 |

- **HER2 Trastuzumab** (Shanehsazzadeh et al., 2023b): 1,240 CDR-engineered variants of trastuzumab targeting HER2 (PDB 1N8Z), with 417 SPR-confirmed binders and 823 non-binders.
- **5A12 VEGF & 5A12 ANG-2** (Minot and Reddy, 2024). Combinatorial CDR mutagenesis of antibody 5A12, validated by yeast display. VEGF: 701 binders and 499 non-binders subsampled from a 642k-variant library (original binder rate 58.4 %). ANG-2: 120 binders and 1,080 non-binders from a 712k-variant library (original binder rate 1.9 %). (PDB 4ZFG/4ZFF)
- **IgDesign Datasets** (Shanehsazzadeh et al., 2023a): Seven therapeutic antigens with SPR-validated binding for designs generated by HCDR3-only or full HCDR123 redesign. Curated set: 1,243 sequences (258 binders, 985 non-binders).

To examine the relationship between GeoFlow-V3’s confidence scores and experimentally measured binding affinities, we construct a non-cherry-picked antibody–antigen subset of SKEMPI 2.0 (Jankauskaitė et al., 2019). First, we identify antibody–antigen complexes by filtering protein descriptions with the keywords *Fab, IgG, mAb, Herceptin, Antibody, scFv, HyHEL-10, and VRC-PG04*. We exclude complexes containing non-canonical amino acids or cases where the annotations do not match the CIF structures. To reduce batch effects and better reflect the goals of *de novo* antibody design, we group mutation data by PDB ID, discard groups with fewer than five entries, and retain only those exhibiting at least a 1,000-fold range in binding affinity. We further remove any samples in which the mutations occur on the antigen chain. These filtering steps yield a high-confidence, antibody-focused SKEMPI subset comprising seven unique PDB complexes (full statistics in Table 2).

**Table 2.** Summary of antibody-antigen subset of Skempi 2.0.

| PDB ID | 1DQJ | 1JRH | 1VFB | 1YY9 | 2BDN | 2JEL | 3HFM |
| --- | --- | --- | --- | --- | --- | --- | --- |
| Count | 32 | 45 | 54 | 20 | 13 | 39 | 85 |
| minimal pKd | 3.5 | 4.5 | 5.1 | 5.9 | 6.5 | 5.5 | 4.7 |
| maximal pKd | 8.5 | 8.4 | 9.2 | 10.3 | 9.8 | 9.0 | 10.5 |
| median pKd | 7.3 | 7.5 | 7.4 | 8.9 | 9.0 | 8.5 | 8.2 |

We benchmark GeoFlow-V3 against several representative baselines.

- We include **AlphaFold Multimer V2.3–Initial Guess**, a variant of AlphaFold Multimer V2.3 (Evans et al., 2021) augmented with the *initial-guess* strategy described by Bennett et al. (2023), where the reference antibody-antigen structure is provided as the initial guess conformation.
- We compare GeoFlow-V3 with our previous model **GeoFlow-V2**, which, like GeoFlow-V3, accepts antibody–antigen sequences together with binding hotspot residues extracted from reference complexes as epitope constraints.
- We evaluate three state-of-the-art all-atom protein structure prediction models: **Protenix-v0.6.0, Boltz-2**, and **Chai-1**.

Following established practice (Bennett et al., 2024; Hitawala and Gray, 2025), we use each model’s ipTM score as a proxy for binding strength. For GeoFlow-V3, we further examine the discriminative power of its internal confidence metrics beyond the ipTM baseline. Specifically, we evaluate:

- **ipTM**, the interface-focused variant of pTM that considers only inter-chain interactions (Evans et al., 2021).
- **ipAE**, the mean predicted alignment error (PAE) for residues in the binder versus the target (Bennett et al., 2023).
- **CDR-ipAE**, the mean PAE restricted to the complementarity-determining regions (CDRs) of the binder versus the target.
- **pLDDT**, the averaged predicted local distance difference test metric from AlphaFold.
- **TA-RMSD**, the target-aligned root-mean-square deviation (RMSD) of the binder.
- **BA-RMSD**, the binder-aligned RMSD of the binder.
- ***GeoFlow-V3 achieves robust performance in binding discrimination***

Binding discrimination performance is quantified using the Area Under the Receiver Operating Curve (AUROC), with results summarized in Figures 5A and 5B. Key observations include:

**Figure 5.**
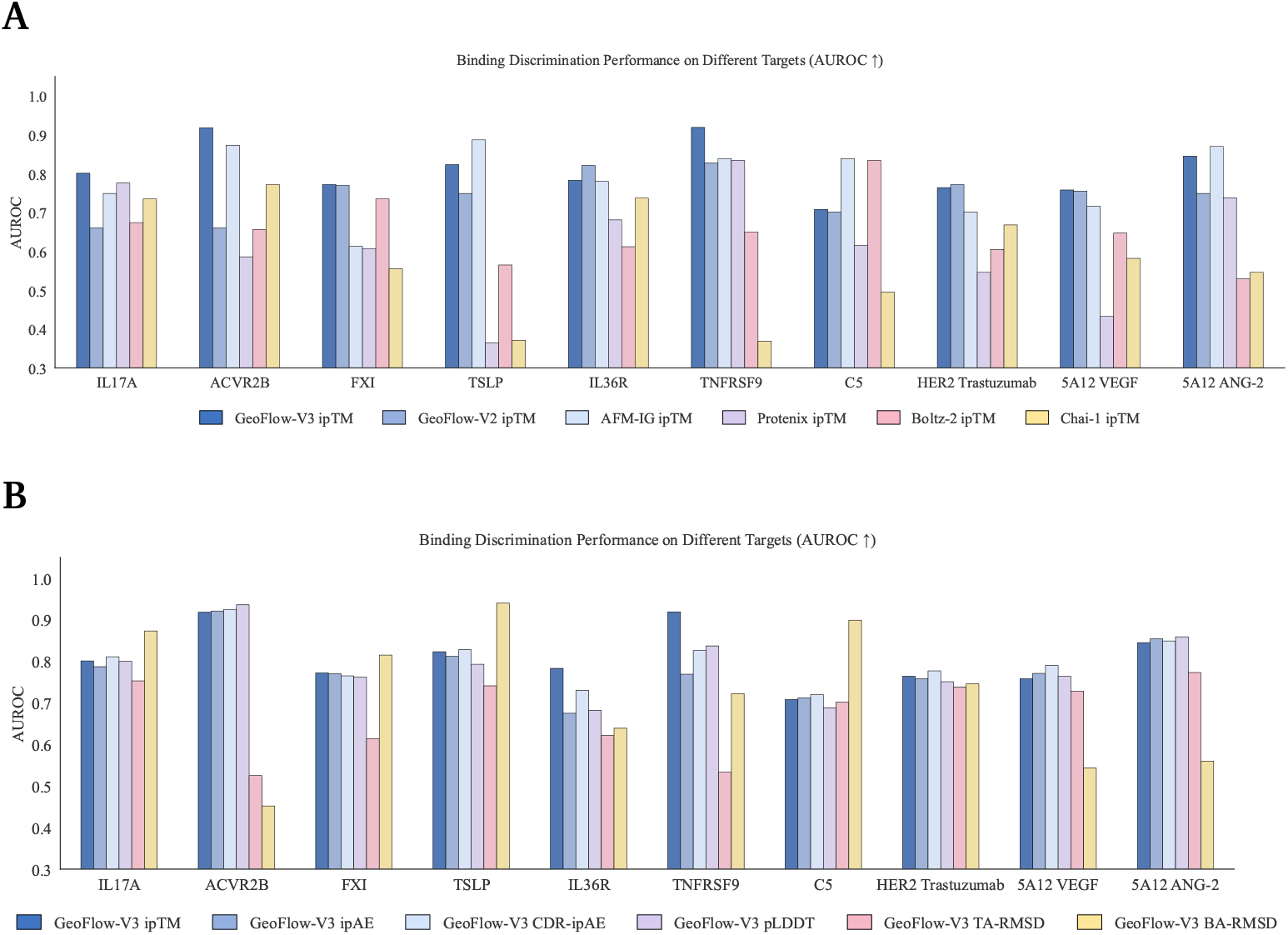
Binder–nonbinder discrimination performance across ten targets, reported as AUROC. **(A)** AUROC comparison of different methods on the binder/non-binder classification task across ten antibody–antigen targets, using each model’s ipTM score as the discriminative feature. **(B)** AUROC of GeoFlow-V3 ‘s own confidence metrics—ipTM, ipAE, CDR-ipAE, pLDDT, TA-RMSD, and BA-RMSD—for the same task, highlighting their relative ability to separate binders from non-binders.

- GeoFlow-V3 delivers improved predictive power over GeoFlow-V2, consistent with the gains in high-DockQ structure prediction and the strong decoy-discrimination capability of its confidence scores (Section 3.1). Antibody–antigen screening is notoriously difficult, particularly due to the challenge of identifying correct docking poses, and we hypothesize that explicit epitope constraints are critical to this performance.
- Surprisingly, AlphaFold Multimer V2.3–Initial Guess surpasses the vanilla diffusion-based all-atom predictors in identifying true binders under most settings. That said, we expect that equipping other structure-prediction baselines with comparable initial-guess or epitope cues could enhance their binding-discrimination capabilities.
- Regarding the performance of GeoFlow-V3’s different confidence metrics, ipTM delivers the most accurate and robust results across all ten targets. BA-RMSD occasionally shows very strong discriminative power but lacks consistency.

#### 3.2.3. Accurate unsupervised binding-affinity discrimination remains challenging

We next examine how well model confidence scores correlate with experimentally measured binding affinities. The ability to accurately rank point mutations is essential for antibody design and optimization. But it is particularly difficult because: (1) antibody–antigen affinity data are scarce and originate from heterogeneous sources with varying antibody format, pH and assay type; and (2) current structure-prediction models are trained on PDB where entities always bind in a complex, making them relatively insensitive to mutations (Agarwal and McShan, 2024; Buel and Walters, 2022), which only slightly changes structural predictions in many cases.

To evaluate this capability, we compute the Spearman rank correlation coefficient between each method’s ipTM scores and experimentally measured affinities in a non-cherry-picked antibody–antigen subset of the SKEMPI dataset. As indicated by Figure 6, discriminating high-affinity binders from weaker ones remains challenging, and no single model consistently dominates across all targets. Some models even show near-zero or negative correlations on specific complexes. Performance across models might be complementary: for example, GeoFlow-V3 performs well on targets 1DQJ, 3HFM, and 2JEL, whereas GeoFlow-V2 shows better performance on 1JRH, 1VFB, and 1YY9. This suggests that ensemble strategies combining multiple models might further improve binding-affinity ranking. Overall, more work needs to be done before we can confidently predict antibody-antigen binding-affinity with a protein foundation model.

**Figure 6.**
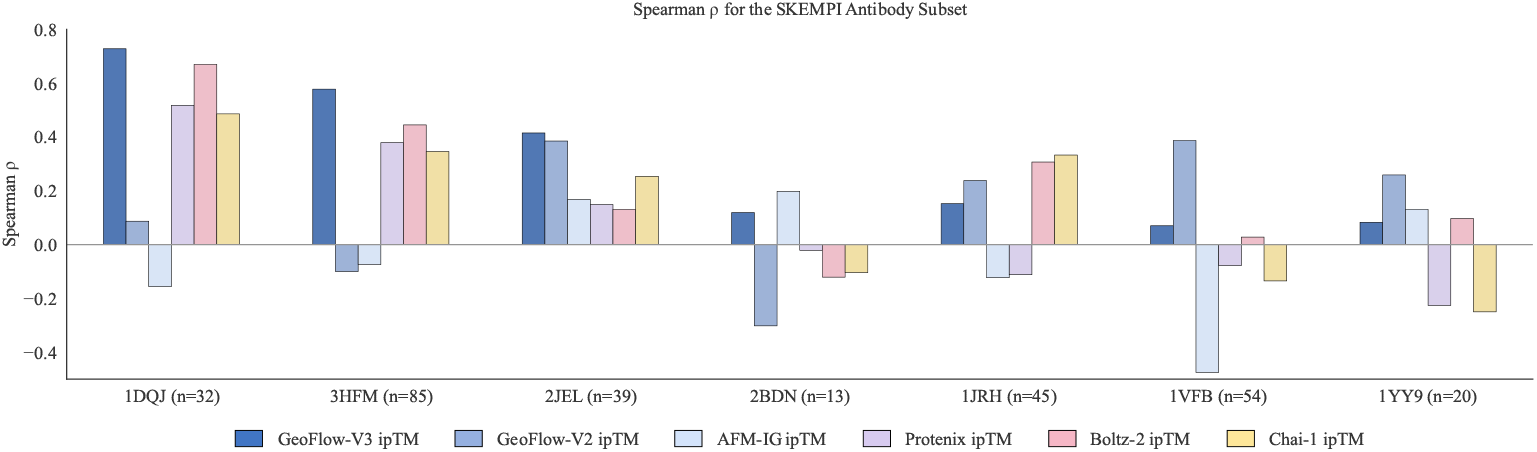
Spearman rank correlations between each model’s ipTM score and experimentally measured binding affinities on the antibody–antigen SKEMPI subset. Despite the difficulty of this task, GeoFlow-V3 shows consistently non-negative, moderate correlations across the seven evaluated complexes, indicating comparatively robust performance.

### 3.3. Case study of *in silico* antibody evolution

We investigate the effectiveness of *in silico* antibody evolution using GeoFlow-V3, with the ipTM score serving as a proxy for binding strength. Case studies focus on five targets: TSLP, IL-33, IL-13, ACVR2B, and IL36R. Nanobodies are designed for the first three targets, which are also experimentally validated (Section 4), while conventional antibodies are designed for the last two as illustrative examples.

For nanobody targets, we adopt the standardized humanized VHH framework h-NbBcII10_FGLA_ (Vincke et al., 2009) as the template and generate 100,000 candidates for both design settings. For antibody targets, the Trastuzumab framework (Cho et al., 2003) is used to generate 10,000 candidates per setting. Designed candidates are filtered *in silico* using thresholds of ipTM *>* 0.8 and target-aligned binder RMSD *<* 3.0. Hexbin plots visualize the filtered designs, highlighting promising candidates that are suitable for downstream wet-lab validation. We also analyze the distribution shift of the Top-100 candidates ranked by ipTM scores to assess how *in silico* evolution improves candidate quality. As shown in Figure 7, *in silico* evolution consistently enhances candidate quality, increasing the number of filtered candidates and improving maximal achievable ipTM scores. While single-round design can produce promising candidates, the evolutionary strategy proves particularly valuable for challenging targets, highlighting its potential as a robust approach for rapid *de novo* antibody design.

**Figure 7.**
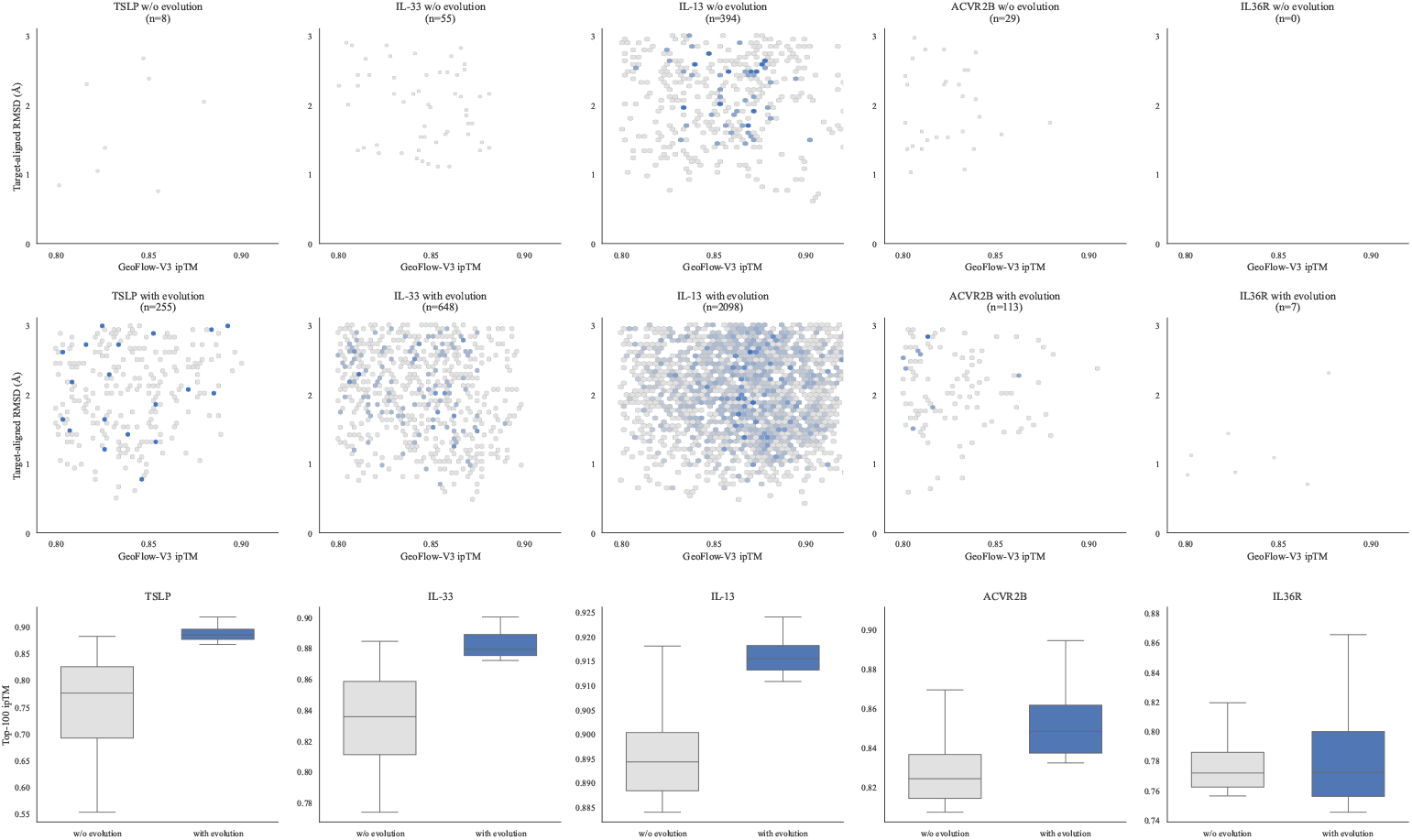
*In silico* antibody evolution strategy enhances candidate quality across five evaluated targets. **First row**: Hexbin plots of promising candidates passing in silico filters (ipTM *>* 0.8 & target-aligned binder RMSD *<* 3.0) without evolution strategy. **Second row**: Hexbin plots of promising candidates passing the same filters with evolution strategy. **Third row**: Box plots depicting the distribution of the Top-100 candidates ranked by ipTM scores for both design settings, with whiskers extending to 1.5 × IQR.

## 4. Rapid *de Novo* VHH Design with Double-Digit Bind Rates

### 4.1. Design protocol

We experimentally validate GeoFlow-V3 on *de novo* nanobody (VHH) design because of this format’s strong therapeutic and diagnostic promise. VHHs are single-domain antigen-binding fragments composed of a single monomeric variable antibody domain. Despite their compact size, nanobodies retain high antigen-binding affinity, can access recessed epitopes, penetrate tissues efficiently, and exhibit exceptional thermal and pH stability. They are also easy to express and purify, making them highly attractive scaffolds for rapid therapeutic development (Harmsen and De Haard, 2007). They have also shown potential to cross the blood–brain barrier under certain conditions (Ruiz-López and Schuhmacher, 2021), further expanding their possible applications. While recent successes in miniprotein binder design are notable (Pacesa et al., 2025; Ren et al., 2025; Zambaldi et al., 2024), *de novo* antibody design remains more challenging, owing in part to the conformational flexibility of complementarity-determining regions (CDRs) and the fact that binding is mediated by loops and *β*-sheet surfaces rather than the *α*-helical motifs typical of mini-binders (Chai-Discovery et al., 2025).

We select five therapeutically relevant targets—**TSLP, IL-33, IL-13, CCR8**, and **PD-1**—to evaluate our *de novo* VHH design pipeline. These proteins are either clinically validated or under active investigation for the treatment of inflammatory diseases and cancers, providing a diverse and medically significant benchmark for evaluation.

- **Cytokines (TSLP, IL-13, IL-33)**. TSLP and IL-13 each activate heterodimeric receptors through two non-overlapping epitopes: TSLP engages TSLPR and IL-7R*α*, while IL-13 binds IL-13R*α*1 and IL-4R*α*. For both cytokines, we design VHH binders targeting the two receptor-binding sites separately, as dual blockade has been shown to inhibit signaling more effectively than single-site inhibition (Savvides et al., 2017). IL-33, which signals through ST2, is similarly targeted at its two spatially distinct ST2-binding regions critical for receptor activation.
- **GPCR (CCR8)**. CCR8, a chemokine receptor implicated in tumor-driven immunosuppression, is targeted at its extracellular loop 2 (ECL2), a key antigenic region.
- **Checkpoint receptor (PD-1)**. PD-1, an inhibitory receptor on T cells central to immune checkpoint therapy, is targeted at its PD-L1 binding interface.

Across these five targets we conduct eight independent VHH design campaigns, including dual-epitope efforts for TSLP, IL-13, and IL-33. Together, these campaigns establish a rigorous benchmark for evaluating the generality, robustness, and performance of our *de novo* nanobody design framework.

For each VHH design campaign, we provide GeoFlow-V3 with up to ten structurally meaningful epitope residues derived from PDB complexes (Table S1) and allow the model to subsample four of them on the fly. This approach is designed to reflect real-world VHH therapeutic development, accounting for two considerations: (1) most targets lack reference nanobody–antigen complexes or even reference nanobody sequences, making precise epitope selection challenging; and (2) epitope subsampling encourages the model to generate diverse binding poses while still targeting the specified region, with the option to narrow down to exact residues if desired. We use the standardized humanized VHH framework h-NbBcII10_FGLA_ (Bennett et al., 2024; Vincke et al., 2009), design all three CDRs with variable lengths, select and experimentally test up to 50 designs per target. We note that the PD-1 design is conducted using a preliminary version of GeoFlow-V3, without the *in silico* evolution mechanism (Section 2.2). For all targets, no known VHH binder—if any exists—is used at any stage of the *in silico* design pipeline.

### 4.2. GeoFlow-V3 achieves high success rate across therapeutically relevant targets

Across all eight VHH design campaigns, at least one binder is identified (Figure 8A & Table S1), demonstrating GeoFlow-V3’s robust capability to generate nanobodies *de novo* for diverse therapeutically relevant targets while respecting user-specified epitopes. The average hit rate reaches 15.5% (excluding PD-1), representing roughly a two-orders-of-magnitude improvement over previous state-of-the-art methods (Bennett et al., 2024) and matching the performance recently reported by Chai-2 (Chai-Discovery et al., 2025). Compared with the preliminary version of GeoFlow-V3, which yields only a single PD-1 binder, the current framework shows a consistent increase in hit rate (though comparison on the same targets would be more direct evidence).

**Figure 8.**
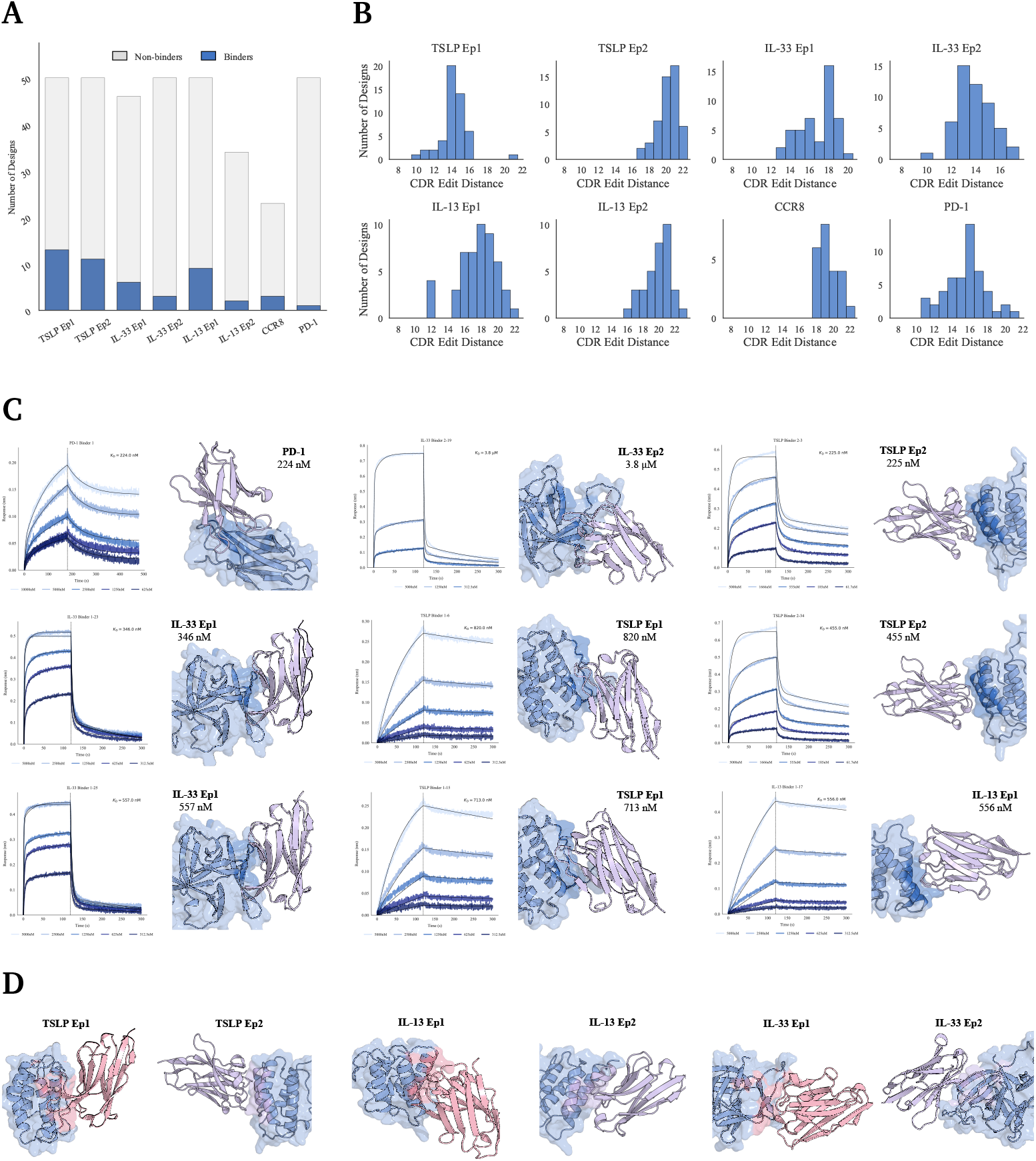
Nanobody design results. **(A)** Success rates for each target, indicating the counts of experimentally validated binding (blue) and non-binding (gray) *de novo* nanobody designs. **(B)** Sequence novelty of experimentally tested designs, measured as the minimum CDR edit distance to known nanobodies in the PDB. **(C)** Representative examples of confirmed binders with multi-concentration BLI curves illustrating binding affinity. **(D)** GeoFlow-V3 generates binders targeting distinct epitopes across different targets.

### 4.3. GeoFlow-V3 designs novel and structurally diverse nanobodies

To assess the novelty of the designed nanobodies, we compute the minimum CDR edit distance between each designed nanobody and all entries in the SAbDab-nano^1^. All SAbDab nanobody sequences are truncated to the Fv fragment and renumbered using the abnumber (Dunbar and Deane, 2016) Python package for consistent CDR annotation. For every designed nanobody, we compute the Levenshtein edit distance for each of the three CDRs relative to every database nanobody. The minimum CDR edit distance for a design is defined as the minimum total CDR distance, i.e., the smallest sum of the three individual CDR distances across all database comparisons. As shown in Figure 8B, all designs have a CDR edit distance of at least 10 from the closest known example, confirming substantial sequence novelty. We further evaluate structural diversity by clustering **confirmed binders** based on antigen-aligned antibody framework RMSD with a 5Å threshold—a metric emphasizing docking pose rather than the intrinsic flexibility of CDR loops. The number of structural clusters for each target, reported in Table S1, reveals that most targets contain multiple distinct clusters, indicating that GeoFlow-V3 can explore several valid conformational states even when epitopes are specified.

### 4.4. GeoFlow-V3 designs nanobodies with substantial affinity

In addition to achieving a double-digit hit rate and generating a diverse binder set, GeoFlow-V3 also yields binders with substantial affinity. Figure 8C presents several confirmed binders and their multi-concentration BLI curves (Section S5.2), showing that GeoFlow-V3 can generate nanomolar VHH binders after only a single round of wet-lab characterization across diverse targets. While there remains room to further enhance binding strength, GeoFlow-V3 eliminates the need for labor-intensive screening campaigns and demonstrates the potential of AI-powered rational antibody design. We further perform binding competition assays of TSLP Ep2 VHH binders against reference anti-TSLP antibodies and evaluate the polyreactivity of confirmed binders toward baculovirus particles (BVP) (Figure S1). These preliminary results demonstrate that GeoFlow-V3 can generate binders with favorable biophysical properties. We are actively improving the pipeline to enhance affinity and to comprehensively assess developability and polyreactivity across designed binders.

## 5. Conclusion, Limitations and Future Directions

We present GeoFlow-V3, a unified atomic diffusion model for both structure prediction and protein design. By combining high-accuracy docking with reliable *in silico* binder discrimination, GeoFlow-V3 eliminates the need for traditional, time-intensive display-based screening. It achieves a double-digit average hit rate in *de novo* nanobody design and can produce sequence-novel nanobodies with nanomolar binding affinity in a single experimental round within three weeks.

Despite these promising results, we acknowledge that GeoFlow-V3 still has important limitations. The current framework does not explicitly optimize therapeutic developability attributes, such as stability, solubility, or polyreactivity, which are crucial for clinical translation. Moreover, although the *in silico* evolution strategy enhances candidate quality, the affinities of identified nanobody binders remain below the threshold typically required for therapeutic use, motivating continued refinement of both the design and screening components.

The demonstrated ability to computationally generate *de novo* antibodies with high success rates underscores a transformative shift toward AI-driven biologics discovery. Owing to its all-atom generative modeling capability, GeoFlow-V3 can naturally extend to a wide range of biomolecular systems, including antigens with post-translational modifications, bispecific antibodies, macrocycles, peptides, and enzymes. Future work will focus on improving affinity throughout the design pipeline, incorporating developability-aware objectives, characterizing therapeutic properties of designed binders, and enabling joint optimization of affinity, specificity, and developability. With these advances, GeoFlow-V3 has the potential to evolve into a versatile platform for rapid, rational, and scalable therapeutic antibody discovery, opening a path toward tackling previously undruggable targets.

## Supplementary information

## S1. Antibody Templates

Following Bennett et al. (2024), we adopt the standardized humanized VHH framework (h-NbBcII10_FGLA_, PDB ID: 3EAK) (Vincke et al., 2009) as the template for nanobody (VHH) design.

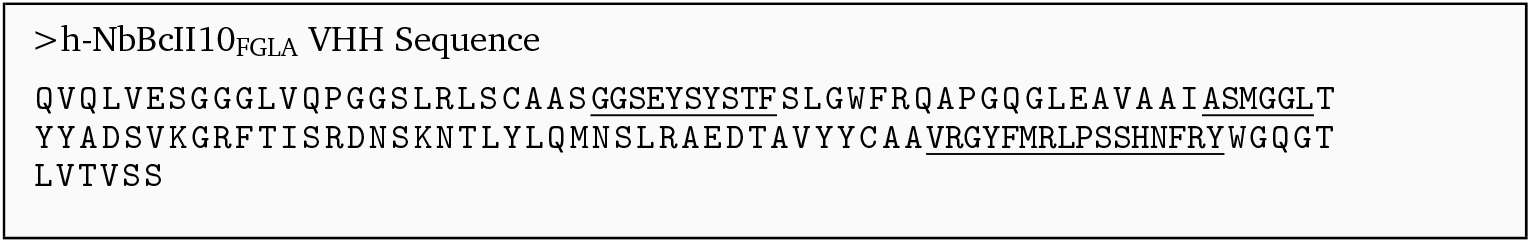

## S2. Target Information

We summarize the target information and their experimental results in Table S1.

**Table S1.** Summary of design targets, per-target design hit rates, and structure clusters.

| Design Target | PDB ID | Binders | Tested | Hit Rate (%) | Structure Clusters |
| --- | --- | --- | --- | --- | --- |
| TSLP (Ep1) | 5J13 | 13 | 50 | 26.0 | 3 |
| TSLP (Ep2) | 5J11 | 11 | 50 | 22.0 | 2 |
| IL-33 (Ep1) | 4KC3 | 6 | 46 | 13.0 | 2 |
| IL-33 (Ep2) | 4KC3 | 3 | 50 | 6.0 | 2 |
| IL-13 (Ep1) | 5L6Y | 9 | 50 | 18.0 | 3 |
| IL-13 (Ep2) | 3BPO | 2 | 34 | 5.9 | 2 |
| CCR8 | 8TLM | 3 | 23 | 13.0 | 2 |
| PD-1 | 8U31 | 1 | 50 | 2.0 | 1 |

## S3. *In silico* Evaluation

We benchmark our method against the following open-source protein-folding models:

- **Chai-1** (Chai-Discovery et al., 2024). https://github.com/chaidiscovery/chai-lab.
- **Protenix-v0.6.0** (Protenix et al., 2025). https://github.com/bytedance/Protenix/tree/v0.6.0.
- **Boltz-2** (Wohlwend et al., 2024). https://github.com/jwohlwend/boltz.
- **AlphaFold Multimer V2.3** (Evans et al., 2021), evaluated using the OpenFold implementation (Ahdritz et al., 2024). https://github.com/aqlaboratory/openfold.
- **AlphaFold Multimer V2.3-Inital Guess**, a variant of AlphaFold Multimer V2.3 incorporating the *initial-guess* strategy described by Bennett et al. (2023). We implement this approach within the OpenFold framework by supplying the reference antibody–antigen complex as a starting template (chain-level) and initializing the structure module accordingly. To prevent information leakage, all complementarity-determining regions (CDRs) are strategically masked.

## S4. Additional Wetlab Results

**Figure S1.**
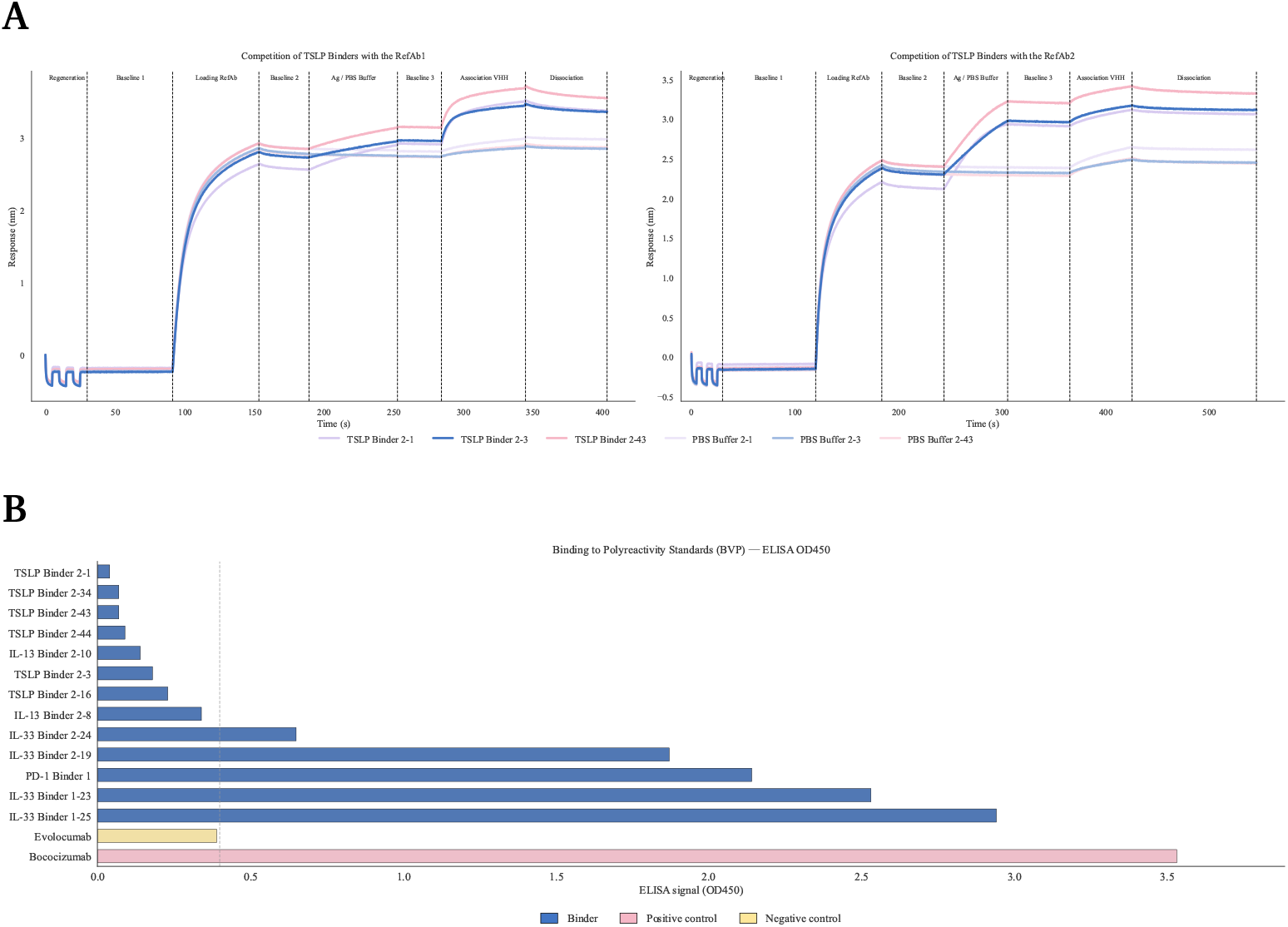
Additional experimental characterization of confirmed binders. **(A)** Binding competition of TSLP Ep2 VHH binders with reference anti-TSLP antibodies RefAb1 and RefAb2. Real-time binding sensorgrams showing sequential loading and association steps for three representative TSLP Ep2 VHH binders compared with the PBS buffer control. **Left**: competition with RefAb1. **Right**: competition with RefAb2. The binders exhibit competition with RefAb2, as evidenced by Association-VHH responses closely matching the PBS control in the right panel, but show no competition with RefAb1, indicating that these VHHs share overlapping epitopes with RefAb2. **(B)** Polyreactivity assessment of confirmed binders against baculovirus particles (BVP). Bococizumab and Evolocumab serve as positive and negative antibody controls, respectively. Nine of the thirteen designs display polyreactivity signals comparable to or lower than the negative control, while the remaining designs show varying levels of polyreactivity risk. These results are consistent with expectations, as the current design framework does not explicitly optimize for reduced polyreactivity, indicating room for future improvement.

## S5. Experimental Methods

### S5.1. Protein production

Genes encoding the selected VHHs were synthesized as gene fragments (Tsingke) and cloned into the pcDNA3.4 plasmid to generate VHH-Fc expression constructs.

Plasmid DNA encoding VHH-Fc was dissolved in sterile water and transfected into HEK293 cells using a transfection reagent. Cells were cultured for 5 days at 37°C, 5% CO_2_, 125 rpm, and 80% humidity. Supernatants were harvested by centrifugation, filtered through a 0.22 µm membrane, and purified via MabSelect SuRe LX resin (Cytiva). Nanobody concentrations were quantified by measuring absorbance at 280 nm (A280) using a NanoDrop spectrophotometer (Thermo Fisher Scientific), and molecular weights were confirmed by SDS-PAGE analysis.

### S5.2. Bio-layer interferometry (BLI)

Binding kinetics were analyzed using the Octet R8 system (Sartorius) with 96-well tilted-bottom microplates. To minimize avidity effects during *K*_D_ determination, monovalent interactions were evaluated by immobilizing VHH-Fc onto ProA biosensors (Sartorius) and using monovalent, Histagged antigens as analytes. Prior to experimentation, ProA biosensors were hydrated in Octet buffer (1× PBS, 0.05% Tween 20) and equilibrated with the sample plate at 30°C for 10 minutes. Biosensors were further equilibrated in Octet buffer for 60 seconds before nanobody loading, with a target loading density set at 2.5 nM.

For the binding hit screen, a one-point BLI assay was conducted using analyte concentrations of either 5 µM or 10 µM. For *K*_D_ determination, a 3- to 7-point series of 2- or 3-fold analyte dilutions was performed. The BLI assay sequence included the following steps: 90 seconds for baseline, 120 seconds for ligand loading, 90 seconds for post-loading baseline, 120 or 180 seconds for the association phase, 180 or 300 seconds for the dissociation phase, and 30 seconds for regeneration. Negative controls, including buffer-only samples, were incorporated for baseline subtraction. Data were globally fit to either a 1:1 or 2:1 binding model across all concentrations to determine *K*_D_.

VHH designs exhibiting a binding-positive curve signature, a signal exceeding 300% of the negative background signal, and a signal greater than 0.1 nm above the background were classified as positive hits. Positive controls with established affinities were included for all targets to validate the assay results.

### S5.3. Binding competition assay

Binding competition between TSLP epitope 2 (Ep2) VHH binders and two reference anti-TSLP antibodies, RefAb1 and RefAb2, was evaluated using the Octet R8 system (Sartorius). RefAb1 and RefAb2 are well-characterized functional antibodies that inhibit the TSLP signaling pathway by binding to two distinct epitopes of TSLP. All experiments were performed at 25 °C in PBST (PBS buffer with 0.005% Tween-20).

The BLI assay sequence included the following steps: 30 seconds for regeneration, 60 or 90 seconds for the initial baseline (baseline1), 60 seconds for RefAb1 or RefAb2 loading, 40 or 60 seconds for the post-loading baseline (baseline2), 60 seconds for TSLP or PBST buffer loading, 30 or 60 seconds for the post-loading baseline (baseline3), 60 seconds for *de novo* TSLP VHH association, and 60 or 120 seconds for the dissociation phase.

Real-time binding responses were measured throughout the experiment. The binding responses of the TSLP VHH binder to TSLP were compared to those of the negative control (PBST), and competitive or non-competitive behavior was determined based on the binding patterns observed.

As shown in Figure S1A, the confirmed TSLP Ep2 binders exhibit competition with RefAb2 but no competition with RefAb1, indicating that these VHHs share overlapping epitopes with RefAb2, aligning with the design expectations.

### S5.4. Baculovirus Particle (BVP) ELISA

The BVP ELISA assay was performed by Sino Biological. Briefly, ELISA plates were coated with 1% baculovirus particle stock (Sino Biological, Lot: 2025.05.21). Next, 100 µL of 150 µg/mL testing VHH-Fc in blocking buffer was added to the wells and incubated for 1 hour at room temperature with shaking. Following incubation, the wells were washed six times with 300 µL of PBS. Each VHH-Fc sample was tested in duplicate. Subsequently, 100 µL of a 0.08 ug/mL dilution of the secondary antibody (Goat Anti-Human IgG (H+L)/HRP) was added to the wells and incubated for 1 hour at room temperature with shaking, followed by an additional six washes with 300 µL of PBS. Finally, 200 µL of TMB substrate was added to each well and incubated for approximately 10 minutes. The reaction was stopped by adding 50 µL of 2N sulfuric acid to each well, and absorbance at 450 nm was measured. For data analysis, Evolocumab (negative control) and Bococizumab (positive control) were also tested. The plotted OD450 values represent the average of two replicates, with the blank subtracted.

Since the molecular weight of Evolocumab and Bococizumab is nearly twice that of VHH-Fc, the molar quantity of VHH-Fc is correspondingly twice as high. Notably, 60% of the designed VHH constructs exhibited OD450 values comparable to or lower than the negative control, highlighting their promising developability as potential therapeutics (Figure S1B).

## Footnotes

1 https://opig.stats.ox.ac.uk/webapps/sabdab-sabpred/sabdab/nano/

